# Confirmation of local transmission of *Orientia tsutsugamushi* during scrub typhus outbreaks in Nepal

**DOI:** 10.1101/852533

**Authors:** Meghnath Dhimal, Shyam Prakhas Dumre, Guna Niddhi Sharma, Pratik Khanal, Kamal Ranabhat, Lalan Prasad Shah, Bibek Kumar Lal, Runa Jha, Bishnu Prasad Upadhyaya, Bhim Acharya, Sanjaya Kumar Shrestha, Silas A. Davidson, Piyada Charoensinphon, Khem B. Karki

## Abstract

**Background:** Scrub typhus is a severely ignored tropical disease and a leading cause of undifferentiated febrile illness worldwide caused by infection of an obligate intracellular bacteria *Orientia tsutsugamushi*. It has been rapidly expanding in South Asian countries, although clear epidemiological information is not available from Nepal. After the 2015 earthquake in Nepal, a sudden upsurge in scrub typhus cases was reported. The objective of this study was to investigate scrub typhus and its causative agents in human, rodent and chigger mites to better understand the ongoing transmission ecology.

**Methods:** Scrub typhus cases with confirmed diagnosis throughout the country were included in the analysis. Studies were concentrated in the Chitwan district, the site of a major outbreak in 2016. Additional country-wide data from 2015 to 2017 was made available from the government database to analyse the disease distribution using geographical mapping.

**Results:** During 2015-2017, 1,239 scrub typhus cases were confirmed with the largest outbreak occurring in 2016 with 831 (67.1%) cases. The remainder 267 cases were reported in 2017. The case fatality rate was 5.7% in 2015 and declined to 1.1% in 2017. Nationwide outbreak of scrub typhus was identified as the cases were found from 52 of the 75 districts of Nepal. A seasonal trend was observed with a peak during August and September (p = 0.01). In addition to the human cases, the presence of *O. tsutsugamushi* was also confirmed in rodents and chigger mites from the outbreak areas of southern Nepal.

**Conclusion:** The detection of *O. tsutsugamushi* in human, rodent, and chigger mites from outbreak locations and wide-spread reports of scrub typhus throughout the country over two years confirms the ongoing transmission of *O. tsutsugamushi* with a firmly established ecology in Nepal. The country’s health system needs to be strengthened for systematic surveillance, early outbreaks detection, and immediate response actions including treatment and preventive measures.

**Author Summary:** Scrub typhus is a disease caused by a bacteria called *Orientia tsutsugamushi* and transmitted to people through bites of infected chiggers (larval mites). After the 2015 Gorkha earthquake in Nepal, a sudden upsurge in scrub typhus cases was reported with repeated outbreaks from different parts of the country. This study has documted epidemiology of scrub typhus and its causative agents in human, rodent and chigger mites confimring the local transmission *O*. *tsutsugamushi* with a firmly established ecology in Nepal. The local transmission of the diseases from most parts of the country demands strengthening for systematic surveillance, early outbreaks detection, and immediate response actions including treatment and preventive measures.

## INTRODUCTION

Scrub typhus is a vector-borne acute febrile illness caused by *Orientia tsutsugamushi* and transmitted to humans and rodents by infected chigger mites (larval stage of Trombiculidae mites) [1, 2]. Historically, scrub typhus was endemic in Asia, Australia and islands in the Indian and Pacific Oceans known as the “tsutsugamushi triangle”[1]. However, there have been recent reports of scrub typhus from Africa, France, the Middle East, and South America suggesting the disease is no longer restricted to this triangle [2]. Scrub typhus is frequently reported from many Asian countries and is endemic in Nepal’s neighboring countries including India (Sub-Himalayan belt) and Bhutan, where it is considered an emerging infectious diseases [3–6]. In Bhutan, one in six undifferentiated febrile patients had rickettsial infections, with scrub typhus being the most common [6]. However, the disease situation in Nepal has remained silent until now, likely due to the high burden of other febrile illnesses with indistinguishable clinical signs and limited proper diagnostic availability.

There have been a few previous attempts to investigate scrub typhus in Nepal. As early as 1981, a study showed the high possibility of scrub typhus in Nepal by showing elevated antibody titers among 10% of healthy adults [7]. Unfortunately, additional surveillance studies in the country were not conducted until 25 years later in 2004. A serological investigation of scrub typhus at Patan Hospital, Kathmandu found a small number of febrile patients (28/876) positive for scrub typhus antibodies [8]. The investigation was performed using a multi-test assay without further confirmation for scrub typhus by immuno-fluorescent assay (IFA), gold standard for scrub typhus diagnosis. It was inconclusive whether scrub typhus or another rickettsial illness, such as murine typhus, was present. Another report in 2007, also indicated the presence of scrub typhus in Nepal [9]. However, no outbreak investigations of scrub typhus (with fatality information) were reported in Nepal before 2014 and no systematic investigations by the government had been conducted. As a consequence, scrub typhus cases had not been reported to the Epidemiology and Disease Control Division (EDCD) of the Ministry of Health before 2014 [10].

In April 2015, Nepal experienced a mega-earthquake claiming thousands of lives, massive destruction, and huge economic losses followed by an upsurge in febrile illnesses [11, 12]. Three months after this devastating earthquake in Nepal (August 2015), a tertiary care teaching hospital in Nepal alerted EDCD that children with fever and severe respiratory features were not responding to the usual course of antibiotic treatment that’s why leading to high mortality (8%) [10, 11]. The usual course of antibiotic treatment was cefexime or ceftraiaxone and imipenem in ICU. After the initially suspected etiologies (Hantavirus and other viral diseases) were ruled out, the samples were screened with /IgM -ELISA for scrub typhus and found positive. This was the first and most significant fetal scrub typhus outbreak in the country [11]. Since then, scrub typhus has been increasingly reported in Nepal but no clear epidemiological picture is available. In this study, a systematic investigation was carried out in Nepal which included patient, vector, and rodent investigations employing both serological and molecular tools.

## MATERIALS AND METHODS

### Ethics statements

This study was approved by the Ethical Review Board of Nepal Health Research Council (NHRC) (Reg. No 305/2016) and conducted in accordance with the Helsinki declarations. All patient records were anonymized prior to analysis. All procedures involved with animal were carried out as per the Ethical Guidelines for the Care and Use of Animals in Health Research in Nepal and followed the regulation of local and national laws regarding the use of animal for research purpose [13].

### Study design and settings

This was a descriptive cross-sectional study conducted in Nepal and included scrub typhus cases of any age and sex reported in 2016 from all the districts throughout the country. Serologically confirmed cases of scrub typhus from various laboratories, including National Public Health Laboratory (NPHL), were reported to EDCD and compiled in the database of EDCD, Ministry of Health and Population (MOHP), Nepal. As a part of outbreak investigation, chigger and rodent studies were also performed from major outbreak area in Chitwan district as detailed in the corresponding section below. Additionally, the aggregated data from 2015 and 2017 were made available from the EDCD submitted by National Public Health Laboratory, and used for mapping. An overall flow-diagram of the study is detailed in Fig 1.

**Figure 1.**
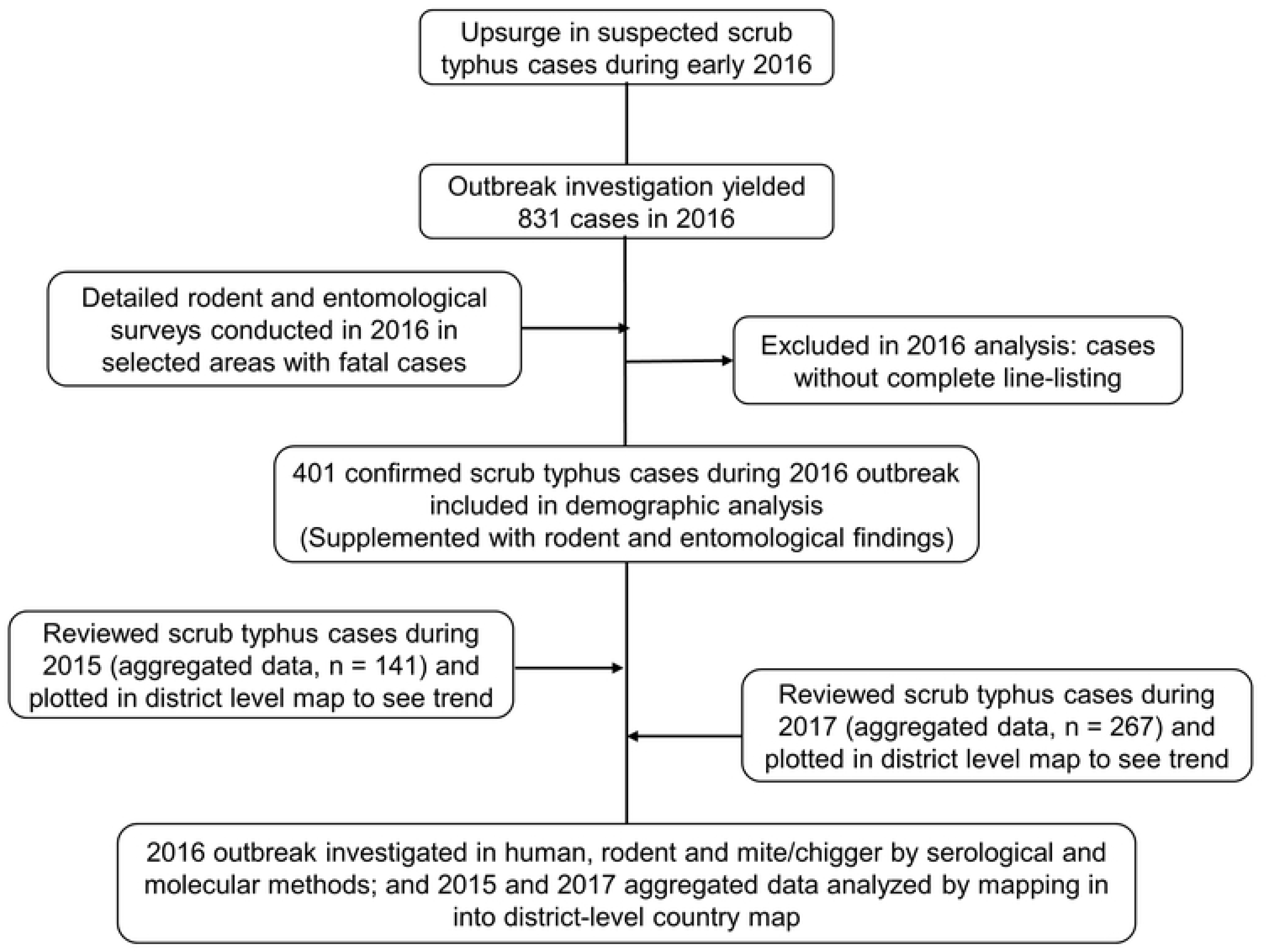
Overall flow-diagramof the study.

### Case definitions

The patients were classified based on the EDCD Guideline on Prevention and Control of scrub typhus adopted from World Health Organization (WHO) [10]. Briefly, 1) a case with acute undifferentiated febrile illness of ≥ 5 days with or without eschar (fever of < 5 days for a case with eschar) was considered “*suspected/clinical case”*; 2) a suspected/ clinical case with an immunoglobulin M (IgM) titer > 1:32 and/or a four-fold increase of titers between acute and convalescent sera was considered as “*probable case”*; and 3) it was declared “*confirmed case* of scrub typhus” when the *O. tsutsugamushi* DNA was detected in eschar or whole blood samples by polymerase chain reaction (PCR), or four fold rise in antibody titers on acute and convalescent sera by IFA which is gold standard assay or Indirect Immuno-Peroxidase (IIP) assay.

### Human sample collection and laboratory investigation for *O. tsutsugamushi* infection

Blood samples were collected from the suspected scrub typhus cases presented to the National Public Health Laboratory (NPHL), Kathmandu and from Chitwan Medical College (CMC) Hospital and Bharatpur Hospital, Chitwan. Samples collected from sentinel sites including CMC, and Bharapur Hospital, Chitwan were transferred to NPHL maintaining cold chain. Samples were tested for scrub typhus using *O. tsutsugamushi* specific IgM-ELISA (Scrub Typhus Detect™ Kit, InBios International, WA, USA) and interpreted as per the manufacturer’s manual. The remaining aliquots of the sample were cryo-preserved at −80°C. Representative scrub typhus IgM positive and negative serum samples (confirmed by IgM-ELISA) were further confirmed by IFA using a panel of *O. tsutsugamushi* specific antigens covered major *O. tsutsugamushi* strains identified in endemic areas at the Armed Forces Research Institute of Medical Sciences (AFRIMS), Bangkok, Thailand and Walter Reed/ AFRIMS Research Unit Nepal (WARUN) as described previously [14, 15]. Briefly, pooled antigens from whole cell-cultured prototype strains including Karp, Kato, and Gillian were used to detect *O. tsutsugamushi* specific antibodies in patient serum samples. Initial screen was performed using dilution of 1:50. Any positive sample was then 2-fold serially diluted to a final concentration of 1:12800 and tested for antibody titer. FITC-conjugated Rabbit Anti-Human IgG; Rabbit Anti-Human IgG-FITC Secondary Antibody was used to determine *O. tsutsugamushi* specific antibodies in humans. Serum titers less than 1:50 were interpreted as negative for IgM and IgG, whereas serum titers 1:50 and less than 1:400 indicated past or recent infection, and serum titer equal to or higher than 1:400 indicated active infection.

### Rodent and entomological investigation of scrub typhus outbreak

#### Site selection

Chitwan district was selected for rodent and entomological surveillance based on the severity and prevalence of the disease in and around this district. The fatality due to scrub typhus mainly occurred in the Mangalpur Village Development Committee (VDC), which is a sub-district level administrative area of Chitwan district. Therefore, three VDCs (Mangalpur, Sharadanagar and Shukranagar) which are located to the South-West of Bharatpur, which is the district headquarters, were selected for the surveillance study (Fig 2).

**Figure 2.**
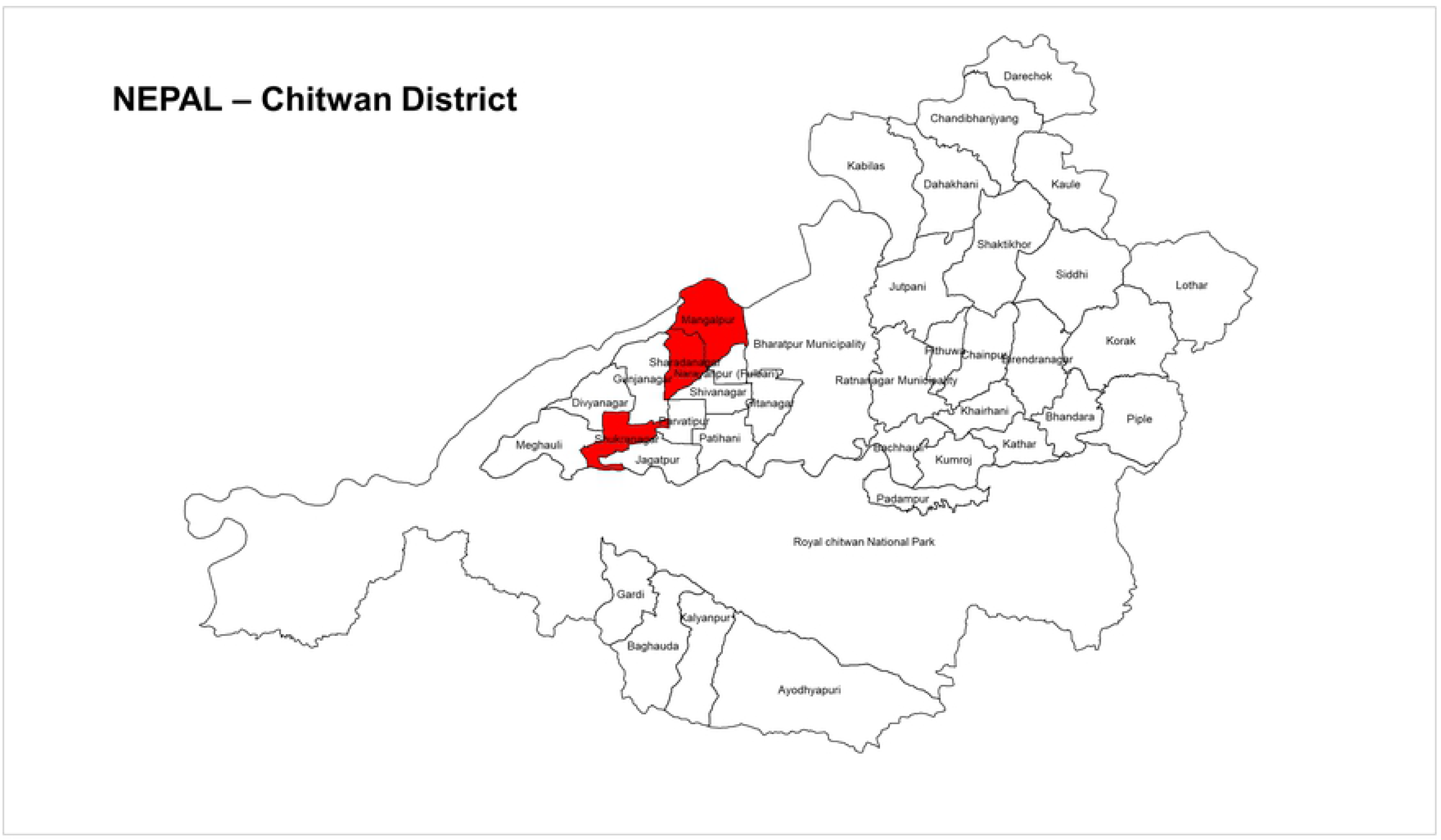
Map of Chitwan district showing the rodentological and entomological study sites. Investigation was carried out in three VDCs (Mangalpur, Sharadanagar and Shukranagar) (dark filled areas)which are located to the South-West of the district headquarters of Chitwan(the outbreak epicenter district). VDC, village development committee (Source: EDCD data using Arc GIS software)

#### Rodent capture

The indicator cases were identified, and rodent trapping was performed in and around the houses of those index cases and their neighbors. Rodents were trapped using Sherman rodent traps tagged with a unique number for identification as described previously [15]. All the traps were equipped with different baits (ripen banana, tomato, or piece of chicken) and placed at various sites (n = 104) inside and outside houses, cattle sheds, near granaries (Aali) and the nearby fields (e.g. kitchen gardens) following the trap lines. Traps were set in the early evening and collected the following morning.

#### Rodent blood collection and species identification

The rodent along with the trap was placed inside a gas chamber and anaesthetized using carbon dioxide inhalation method (2-3 L/ h). Following the procedure of euthanasia, the rodent was taken from the trap, weighed, and essential characteristics were recorded for species identification. Blood was collected by direct cardiac puncture using 3 mL disposable syringes. Serum samples pipetted into cryo-tubes were kept in a cold box with pre-frozen ice packs at −80°C and immediately shipped to the laboratory of Walter Reed/AFRIMS Research Unit Nepal (WARUN) at Bharatpur Hospital and then to WARUN in Kathmandu in cold box maintaining temperature at −80 °C. The samples were analyzed at WARUN for further related investigations. Rodents were identified to species according to their length, color patterns, tail, pinna, number of mammary glands and color of incisor teeth.

### Mite collection, identification and processing

Mites and other ectoparasites were collected by combing the anaesthetized rodent’s outer surfaces (ventral and dorsal surfaces, arm-pits, groins, and areas near pinnas, etc.) onto a white clean paper sheet to help identify the ectoparasites. The ear pinna of rodents was examined under a dissecting microscope for chiggers and individual or clusters of chiggers were removed together with thin layer of ear skin using fine forceps. Chiggers were transferred into labelled vials containing 70% ethanol and stored in an ice-cold box for preservation. Representative chiggers from each rodent host were randomly sampled for morphological identification. Permanent slides of chigger samples were prepared and mounted at WARUN laboratory in Kathmandu, Nepal following method described previously (reference: https://www.ncbi.nlm.nih.gov/pubmed/30282787).

### Laboratory investigation of non-human samples (rodents and chiggers)

#### *O. tsutsugamushi* specific antibody detection in rodent serum by IFA assay

The rodent serum samples were tested for the presence of IgG antibody against *O. tsutsugamushi* by IFA assay with a panel of *O. tsutsugamushi* antigens described previously for the human samples. The seropositivity cut off value for interpreting infection in rodent sera for IgG was 1:50.

#### *O. tsutsugamushi* specific PCR using rodent tissue samples

Lung and liver tissues from rodents were used for pathogen detection. Tissue samples were dissected and homogenized in buffer ATL (Qiagen, Hilden, Germany) with sterile 5 mm stainless steel beads and processed using Tissuelyzer LT (Qiagen). The homogenate was lyzed with proteinase K and subjected to DNA extraction using the QIAmp DNA mini kit (Qiagen). Subsequent quantitative polymerase chain reaction (qPCR) amplification for *O. tsutsugamushi* was performed using specific primer targeting 47-kDa membrane protease protein encoding gene (htrA) as described previously [15]. Positive samples were confirmed by amplification of 56-kDa tissue specific antigen gene (TSA) and the DNA fragment was resolved using a 1% agarose gel electrophoresis and UV system.

#### *O. tsutsugamushi* specific PCR using chigger samples

Each chigger mite was placed in a tiny drop of phosphate buffered saline (PBS, pH 7.4) under a dissecting microscope. The exoskeleton and internal tissue contents were separated by a puncture and squeeze method. The chigger exoskeleton was mounted on a slide for species identification, while the internal contents was homogenized and used for PCR identification of *O. tsutsugamushi* using the 56-kDa-TSA gene as described above for rodent tissues.

### Data analysis

Data were entered in Microsoft excel and analyzed in IBM SPSS version 22.0. Data were analyzed by chi-square for proportions as applicable. To understand the countrywide situation of scrub typhus, cases were plotted into the district-level country maps.

## RESULTS

### Socio-demographic findings

In 2016, a total of 831 cases with 14 deaths (Case fatality rate (CFR) = 1.7%) were reported from 47 districts throughout the country (Fig 3). Although 831 cases were reported in the country during April-December 2016 in Nepal, complete line listing was available for only 401 cases (male = 163; female = 238 from EDCD. The mean age (years) ± standard of the cases was 28.9 ± 18.2 years and median was 25.0 (IQR14.0-42.0) years. More than half of the cases (57.1%) were below 30 years of age, of that 14.5% were children below 10 years of age (Table 1). The majority of the cases belonged to Janajati/Aadhibasi (44.4%) and Brahmin/ Chhetri (44.1%) ethnic groups with no gender-based differences.

**Figure 3.**
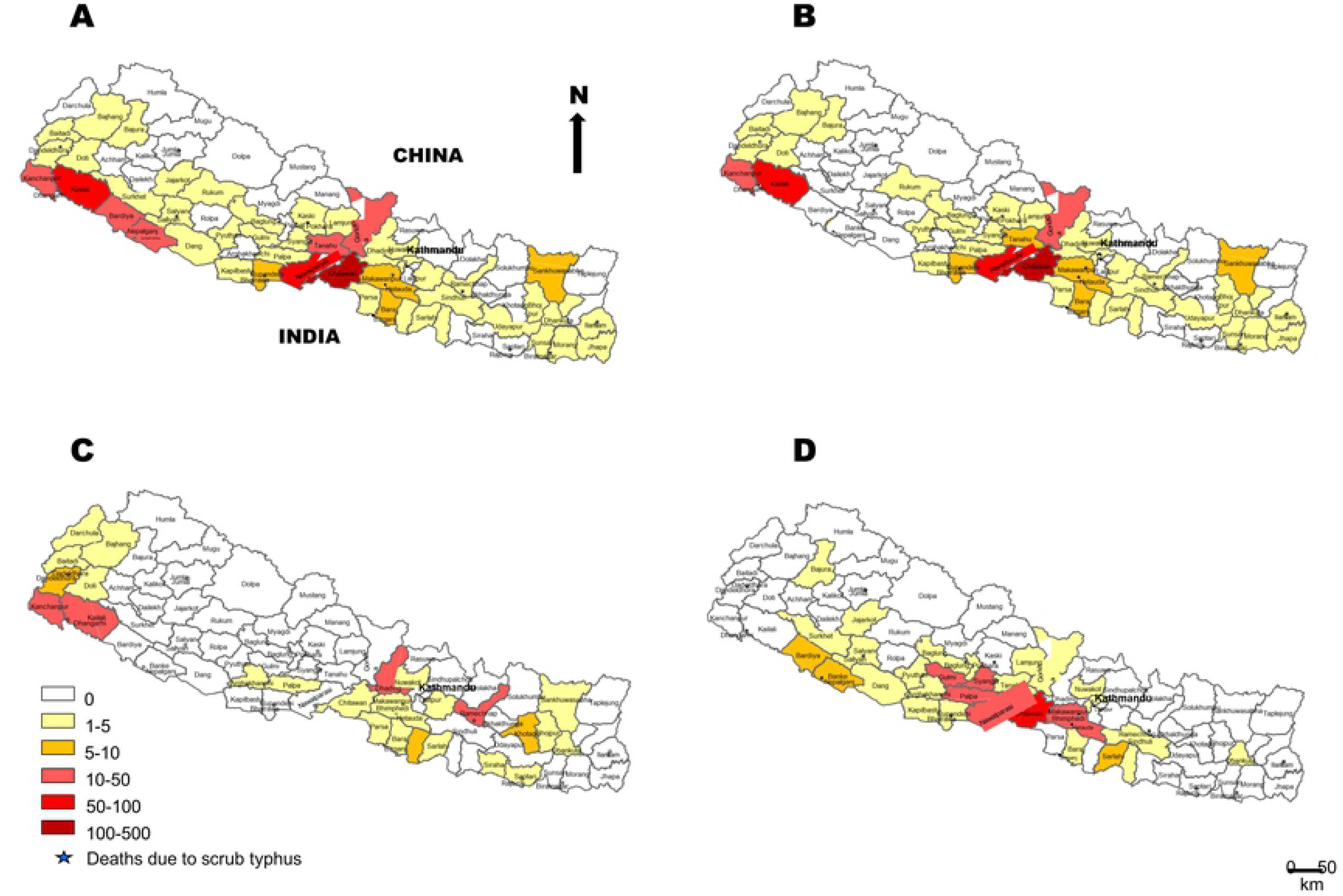
Distribution scrub typhus cases during major outbreaks in Nepal. In 2016, a total of 831 scrub typhus cases (based on peripheral reporting) were reported from 47 of the 75 districts (A), of which a total of 401 cases from 42 districts were confirmed at national reference laboratory and included in further analysis (B), while 141 scrub typhus cases from 25 districts were reported to EDCD in 2015 based on aggregated data available from EDCD (C), and a total of 267 cases were confirmed from 30 districts in 2017 (D). Respectively 8, 14 and 3 deaths due to scrub typhus were confirmed in 2015, 2016 and 2017. From 2015 through 2017, 52 out of 75 districts of the country reported cases of and/or deaths due to scrub typhus (Source: EDCD data using Arc GIS software).

**Table 1:** Scrub typhus cases diagnosed in Nepal by health facilities, 2016

| **Health facility/ area** | **Number of cases (%)** |
| --- | --- |
| Chitwan Medical College, Chitwan | 509 (61.3) |
| National Public Health Laboratory, Kathmandu | 252 (30.3) |
| Bharatpur Hospital, Chitwan | 45 (5.4) |
| Narayani Sub-regional Hospital, Birgunj | 16 (1.9) |
| Siddhartha Hospital, Kailali | 9 (1.1) |
| Total | 831 (100.0) |

Overall, young adults and children were the majority affected by scrub typhus in Nepal, and this proportion was significantly different among age groups (*p* < 0.001) (Table 2). Among male scrub typhus patients, the age skewed towards children (22.8%) while among females, cases were more in age group 20-29 years (27.1%). There was no difference in the median age of scrub typhus patients by gender (*p* = 0.237) (Table 2)

**Table 2:**
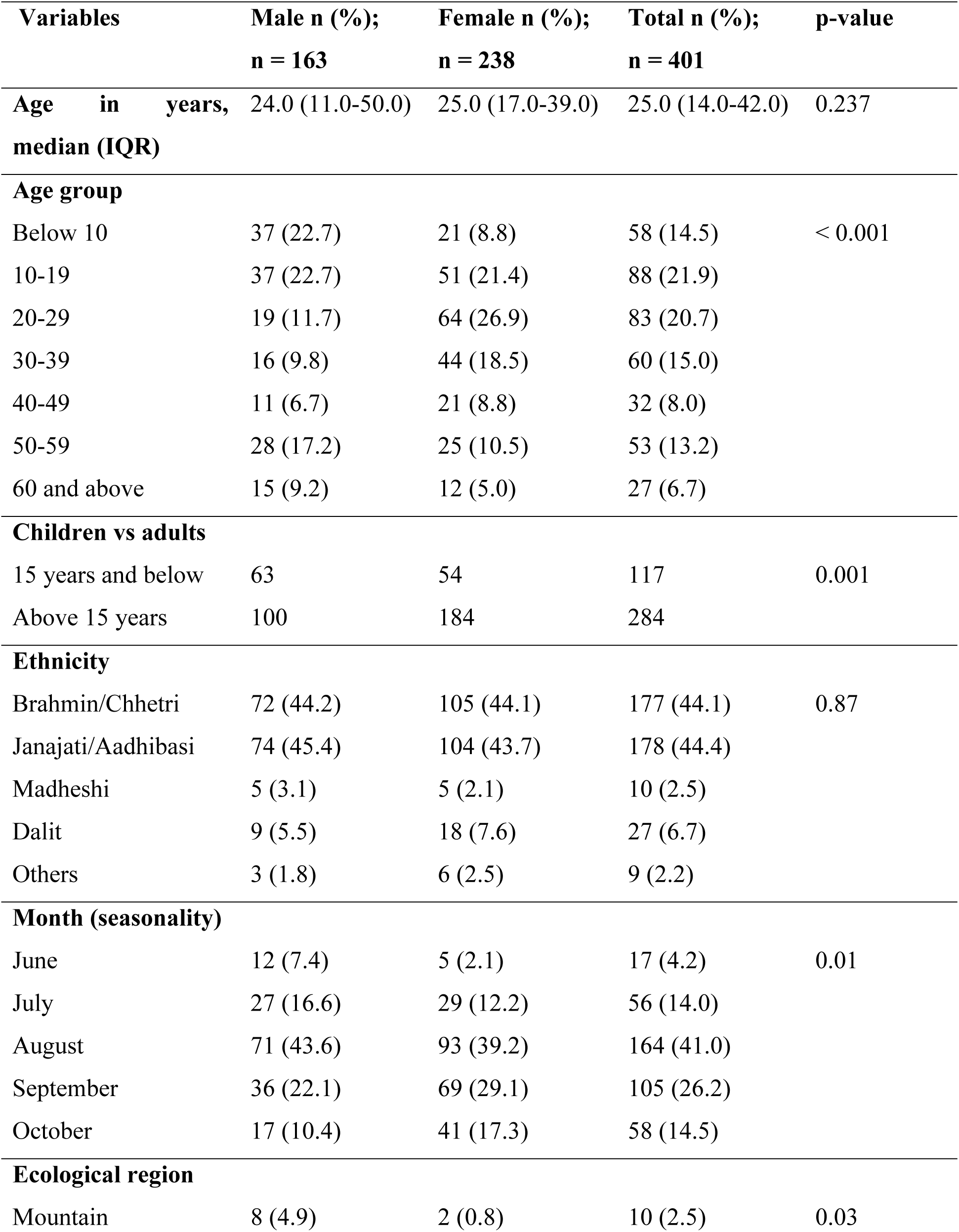
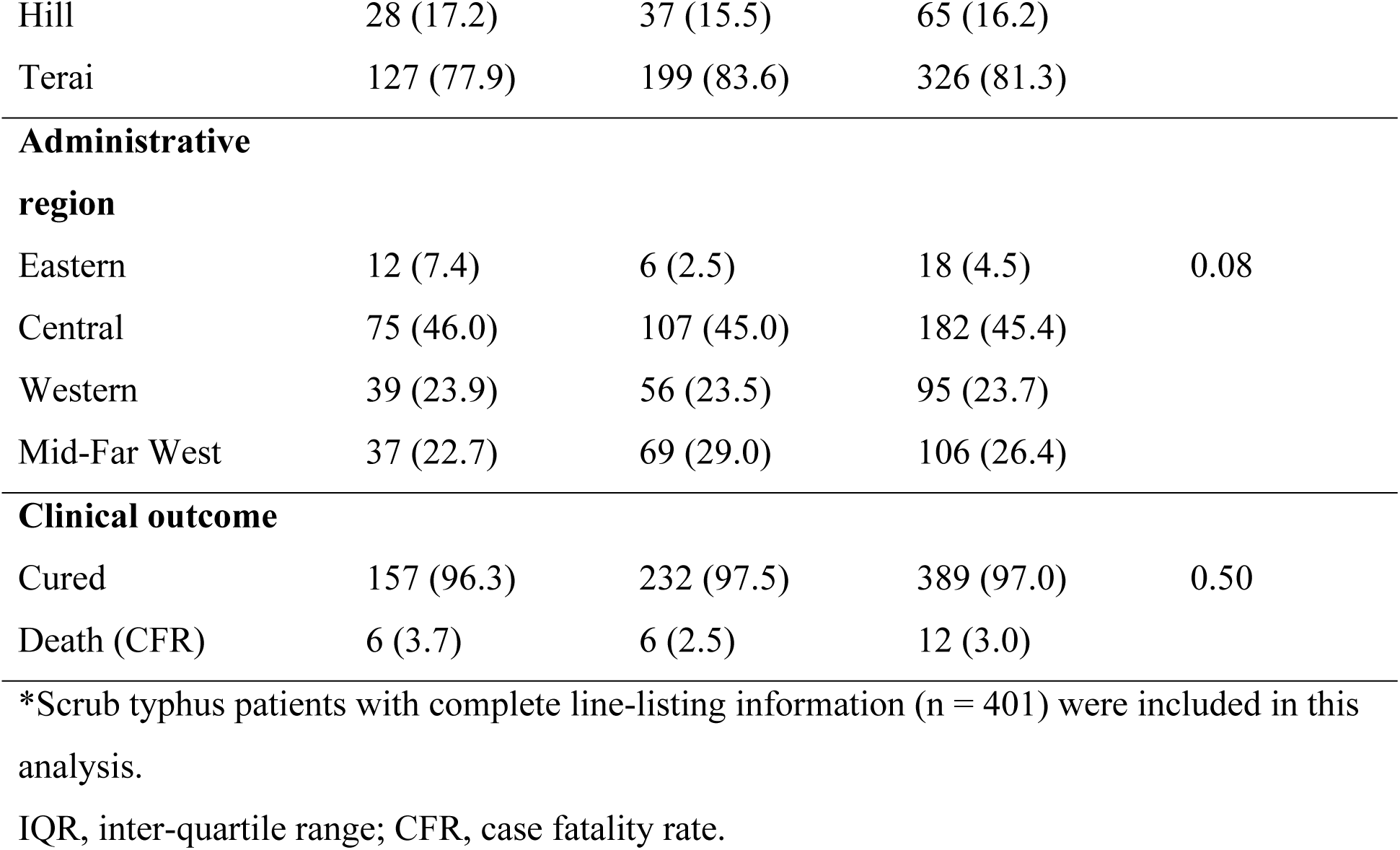
Epidemiological characteristics of scrub typhus cases in Nepal, 2016*

### Geographical and temporal distribution of scrub typhus in Nepal, 2016

The vast majority of scrub typhus cases were from Tarai (81.5%) (*p* = 0.03), the low-land ecology of southern Nepal bordering northern India and the regional analysis identified the central region (45.6%) as the most affected area in the country (Table 2, Fig 3). The most affected district was Chitwan contributing to 34.4% of the total cases (n = 138), followed by Kailali (n = 63), Nawalparasi (n = 55), Kanchanpur (n = 26), Gorkha (n = 15) and Tanahun (n = 10) (Fig 3, Suppl Table 1). The 2016 outbreak of scrub typhus demonstrated a seasonal trend in Nepal that peaked in the month of August (n = 164; 40.9%) and September (n = 105; 26.2%) (*p* = 0.01) (Fig 4, Table 2).

**Figure 4:**
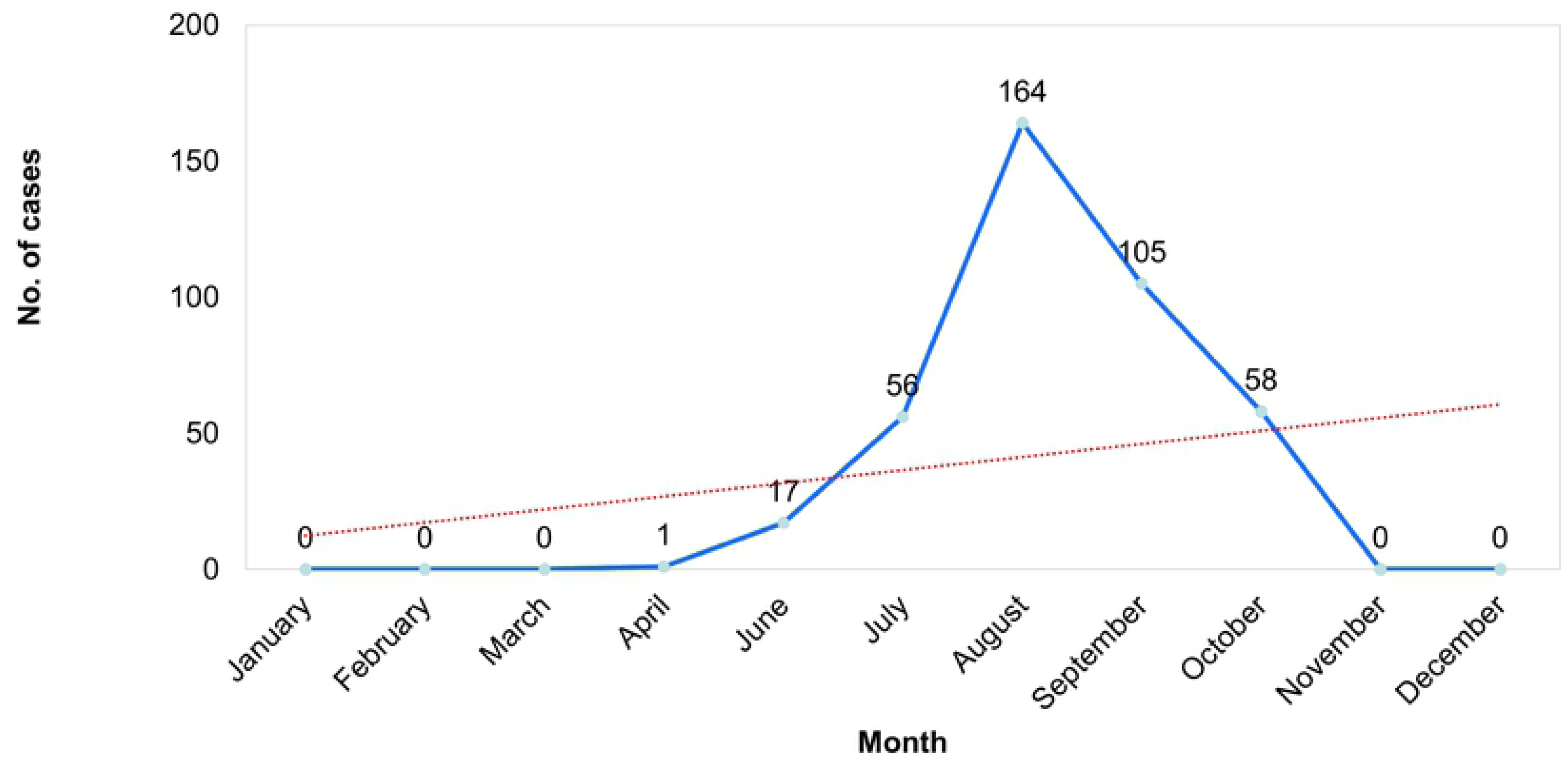
Seasonal trend of scrub typhus in Nepal, 2016.

### Diagnosis, outcome and trend of scrub typhus cases in Nepal, 2015-2017

In 2016, cases were reported or confirmed at five health facilities (Table 1) mostly from the outbreak epicenter, Chitwan district (61.3%). Representative serum samples (n = 61) tested by IFA further confirmed the presence of that *O. tsutsugamushi* infection in Nepal (Table 3). From the 33 samples which tested positive by ELISA, 28 also tested positive by IFA. Similarly, one out of 28 ELISA negative sample was positive by IFA. Regarding the prognosis of the disease, 12 cases (3%) (6 males and 6 females) died while the rest were cured after treatment. After analyzing the cases reported to EDCD during 2015-2017 (n = 1239), it was determined the disease was firmly established in Nepal and a large outbreak occurred in 2016 (n = 831) involving 47 districts. An additional 267 cases were reported in 2017 and there were 141 in 2015 (Table 4). Twenty-five people died due to scrub typhus during 2015-2017 with an overall CFR of 2.0, decreasing from 5.7 in 2015 to 1.1 in 2017 (Table 4). The district wise reports of Scrub typhus from 2016 to 2017 is presented in Supplementart table (Table S1).

**Table 3:** Laboratory findings of selected human samples underwent IFA confirmation

| Sample collection sites | Total samples | Positive by<br>RDT (%) | Positive by IgM<br>ELISA (%) | Positive by<br>IFA (%) |
| --- | --- | --- | --- | --- |
| NPHL | 50 | NT | 30 (60.0) | 26 (52.0) |
| Bharatpur hospital | 11 | 3 (27.3) | NT | 3 (27.3) |

A fraction of samples (n = 61) from suspected scrub typhus cases were randomly selected and sent to reference laboratory (Armed Forces Research Institute of Medical Sciences, Bangkok, Thailand) for IFA confirmation. RDT, rapid diagnostic test; ELISA, enzyme linked immune-sorbent assay; IFA, indirect fluorescence antibody assay; NPHL, National Public Health Laboratory; NT, not tested.

**Table 4:** Trend of scrub typhus cases and deaths in Nepal, 2015-2017

| Year | No. of<br>districts | No. of cases<br>(%) | No. of deaths<br>(%) | Case fatality<br>rate, % |
| --- | --- | --- | --- | --- |
| 2015 | 25 | 141 (11.4) | 8 (32.0) | 5.7 |
| 2016 | 47 | 831 (67.1) | 14 (56.0) | 1.7 |
| 2017 | 30 | 267 (21.5) | 3 (12.0) | 1.1 |
| Cumulative, 2015-2017 | 52 | 1239 (100) | 25 (100) | 2.0 |

### Rodentological and entomological investigation identified hosts of *O. tsutsugamushi* in the outbreak areas of Nepal

Of the total 104 traps used, 12 small rodents were successfully trapped and 9 of 12 were identified as member of *Rattus rattus* species complex, while 3 remaining rodents were *Suncus murinus.* (Table 5). Out of 12 rodents, three (25%) had chigger mite infestation. Three rodents each had 4, 3 and 3 mites, respectively. The chigger index was 0.92. Details of rodents and chiggers have been presented in Tables 5 and 6.

**Table 5:**
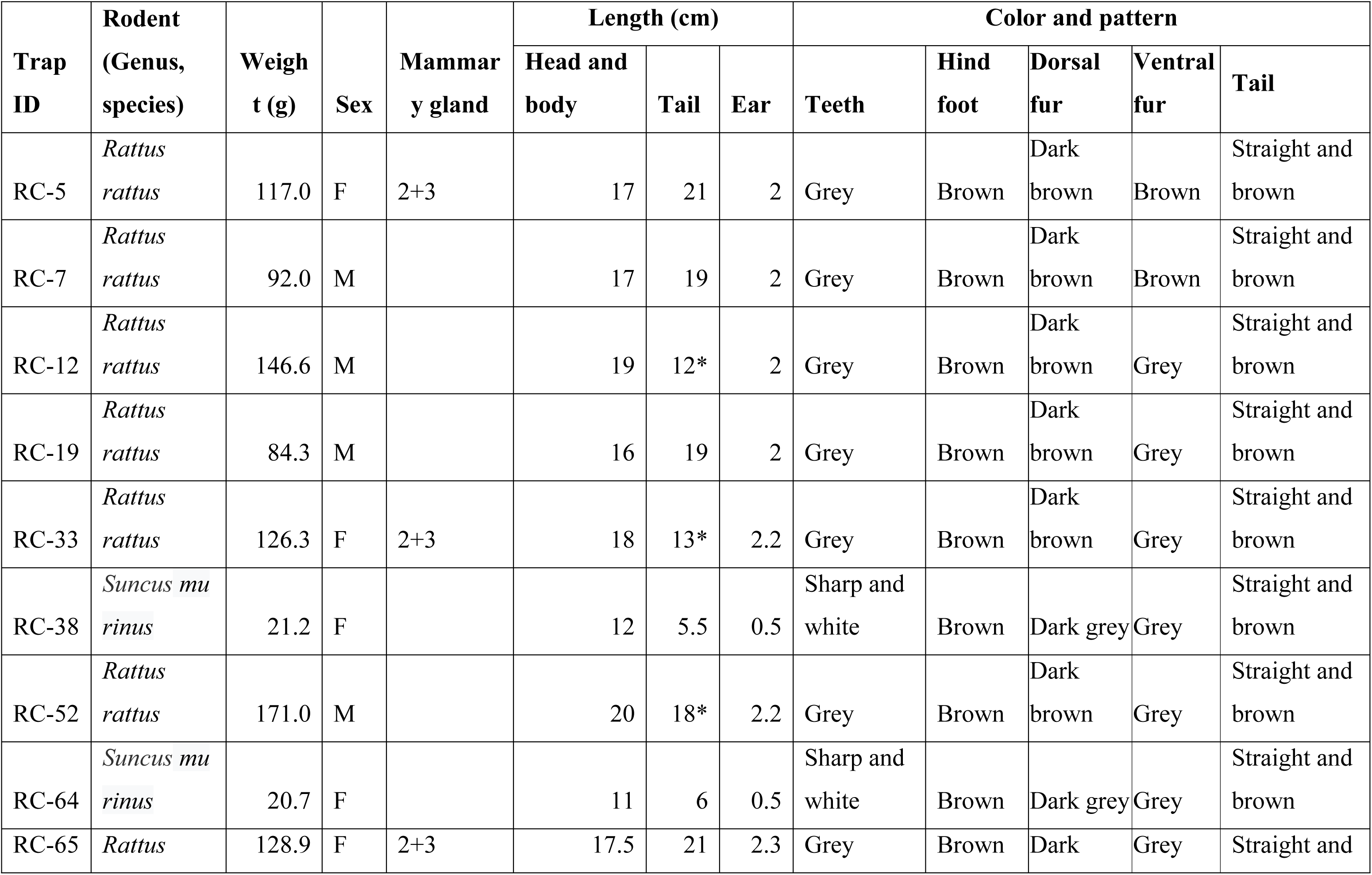
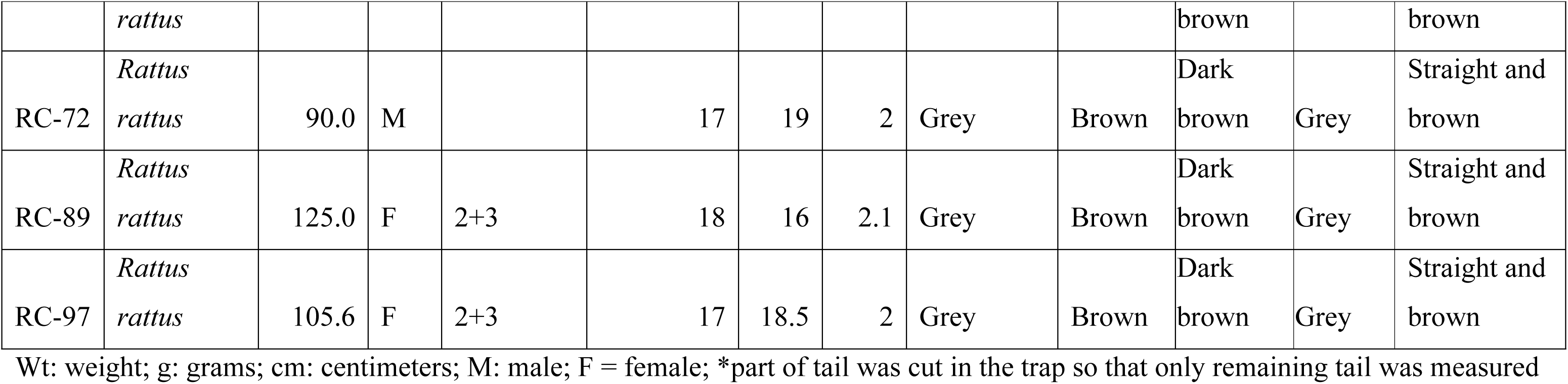
Characteristic features of rodents captured during scrub typhus outbreak in Nepal

**Table 6:**
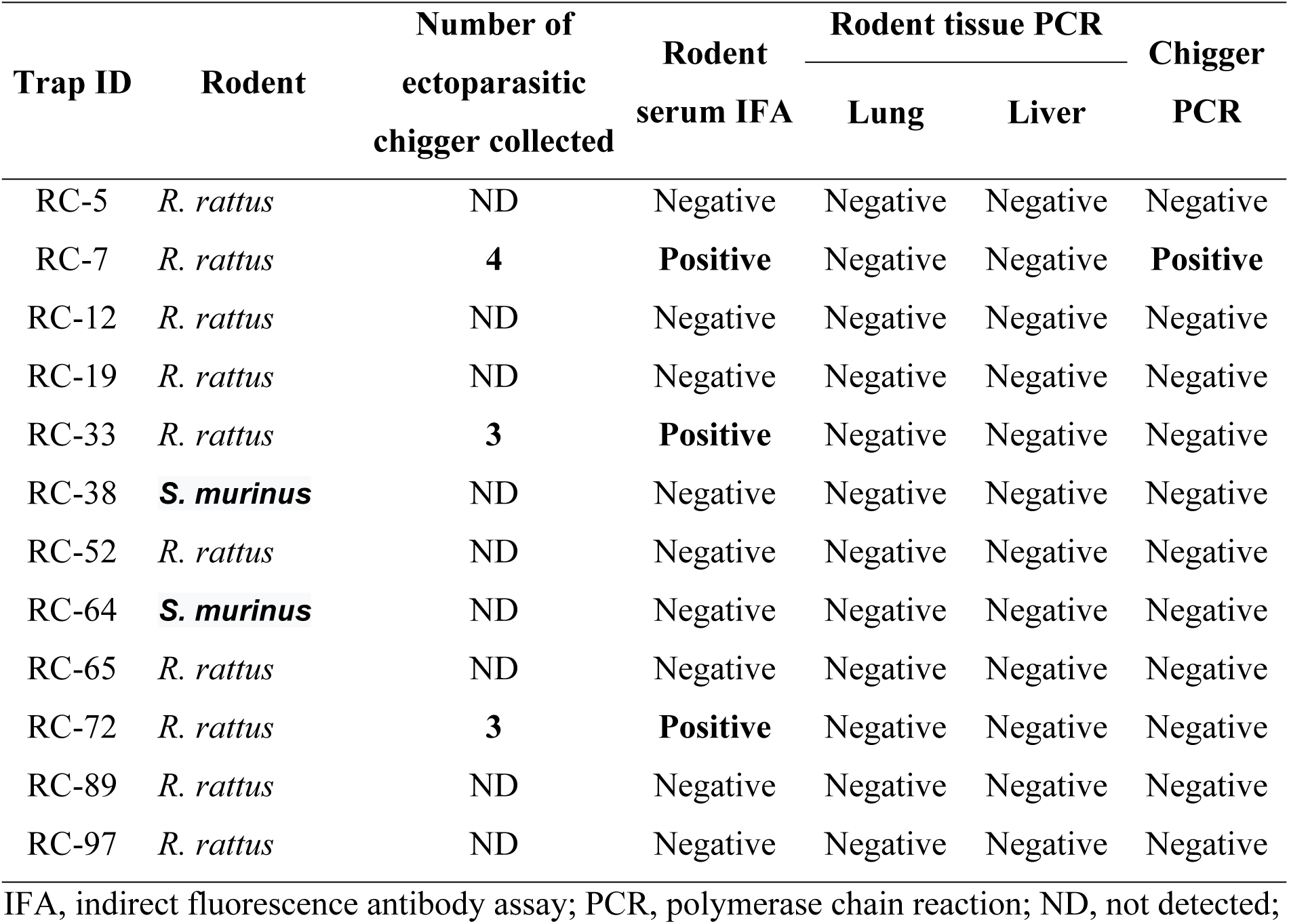
Serological (IFA) and molecular findings of rodents and chigger samples during scrub typhus outbreak in Nepal

### Confirmation of *O. tsutsugamushi* infection in rodents and chiggers collected in the outbreak areas of Nepal by IFA/ molecular techniques

Three rodent serum samples out of nine were confirmed scrub typhus positive by IFA (IgG titer > 50) (Table 6). However, all the 24 rodent tissue samples (12 lungs and 12 livers) were found *O. tsutsugamushi* negative by PCR, suggesting that no active infection was observed in this rodent population. Similarly, one of the three chigger samples was also confirmed *O. tsutsugamushi* positive by PCR.

## DISCUSSION

Our study shows scrub typhus as an emerging public health problem in Nepal with several outbreaks since 2015. Scrub typhus is a severely understudied neglected tropical disease and a leading cause of undifferentiated treatable fever in Asia [1, 16]. This study uncovered a firmly established nature of scrub typhus outbreaks in Nepal through evidences of the causative agent *O. tsutsugamushi* in human (patients), animals (rodents) and vector/ reservoir hosts (chigger mites) during the recent fatal outbreaks. Despite the frequent reports of outbreaks from neighbouring countries, particularly India and others [3-6, 17-20], scrub typhus remained silent in Nepal for decades after its first indication in 1981 [7].

This has resulted in a significant lack of understanding when it comes to epidemiological features, treatment response and severity, and its ecological niche. This type of baseline information is required for a country to formulate appropriate guidelines and strategies for scrub typhus control. Apart from a handful reports discussing the human cases [7-10, 21, 22], no study explored whether the human-host-vector/ reservoir-pathogen cycle is maintained in the country. This ecological chain is essential for an abrupt outbreak of any magnitude [23]. The present study has contributed by providing solid evidence of the ongoing circulation of the scrub typhus pathogen *O. tsutsugamushi* among human, rodents, and chigger mites in areas affected by recent large outbreaks.

The scrub typhus outbreaks that occurred in Nepal during 2015-2017 may be linked to the devastating earthquake of 2015. The outbreaks could have been triggered as a result of intimate contact between human beings and rats that might have come out of their usual underground habitat with the demolition of many houses [24]. Close proximity while living in temporary shelters [12] due to overcrowding and unsanitary conditions could have contributed to increased contact between vectors, pathogens and humans [24]. A weakened health system due to the massive earthquake compromised the availability of diagnostic and treatment facilities which further affected the control program resulting in the large outbreak in 2016. Without the massive earthquake, the likely boosted for bursting out this fatal outbreak of scrub typhus in the country.

A largescale scrub typhus outbreak during 2015-2017 initiated with an abrupt outbreak in 2015 with 141 cases and 8 deaths, which was an unforeseen eruption of this disease after the years of silence in the country. A complete disappearance of the disease in a territory for a long period before a sudden re-emergence in an epidemic form has remained phenomenal in scrub typhus. For example, scrub typhus re-emerged in Maldives as a fatal epidemic in 2002-2003 after 58 years of latency [25, 26], and it also resembles the scenario in India where the re-emergence observed during 1990s after World War II [4]. There was no strong evidence of persistent scrub typhus in the Nepal after the initial indication in 1981, although we cannot totally exclude this possibility considering the lack of diagnostic facilities, endemicity, and inadequate clinical suspicion/precision in Nepal. A common misdiagnosis as typhoid fever based on highly cross-reactive Widal test in Nepal [12] and the minimal clinical interest (due to effective treatment) would also be other factors [25]. Nevertheless, we cannot overlook the serologically positive cases reported in 2004 and 2007 in Kathmandu valley [8, 9] when considering this latency.

Among those infected with scrub typhus, the majority were females < 40 years old involving a significant proportion of children during the investigation. Younger and reproductive group of female in countryside are mostly involved in outdoor or agriculture activity in Nepal, could be the possible reason for increased infection with scrub typhus. India, South Korea and China also reported higher incidence among female [17, 27, 28]. Similarly, we observed a clear seasonality with the majority of cases being detected in August and September, which is quite similar to what reported from the Indian states [17, 20]. The overall pattern of scrub typhus in Nepal resembles to neighbouring countries indicating the potential cross-border transmission.

In Nepal, scrub typhus cases were reported nationwide (52 of 75 districts) in just three years since the disease may expand very rapidly as seen in other parts of the world [29, 30]. In this study, the majority of cases were reported from lowland terai districts where other febrile illnesses including Japanese encephalitis (JE) [31, 32], leptospirosis [33], and dengue [31, 32] had been frequently reported. Even before the initiation of JE vaccination in Nepal, only one-third of the acute encephalitis syndrome (AES) cases were due to JE, which declined with immunization, however the AES still remains persistent clearly indicating other aetiologies of AES in the country. From the same areas of Nepal, cases of leptospirosis and dengue were identified among AES population [32, 33]. Looking at the severity and high fatality (up to 6%) of scrub typhus during 2015-2017, it is logical to consider that a fraction of AES cases could be due to scrub typhus and vice-versa in Nepal. This is further supported by the contemporaneous outbreaks occurred in some states of India (along the Nepal border) where scrub typhus was identified as one of the significant causes of AES [19, 23]. Interestingly, 6 of 8 fatal cases reported in that Indian state had evidence of *O. tsutsugamushi* infection [19]. In addition to AES cases, rodents and mites were also found positive for *O. tsutsugamushi* in those AES-reported areas [23]. This is very much similar to what we found in Nepal suggesting the need to include scrub typhus in the differential diagnosis in AES and other febrile illness in Nepal.

Despite the high fatality (6%) observed during the first wave of scrub typhus in Nepal in 2015, CFR successfully declined to approximately 1% in 2017 which is quite encouraging for the disease control program. This rate is within the wide range of reported fatality (median, 6% and 1.4% for untreated and treated cases, respectively) [1]. The government initiatives that perhaps lowered down the CFR [10, 11] include distribution of guidelines, public awareness, and training and orientation of physicians/ health workers for prompt treatment of scrub typhus with the available drugs (doxycycline, azithromycin, chloramphenicol and ciprofloxacin or their appropriate combinations). Infection with resistant or reduced drug-susceptible *O. tsutsugamushi* strains often yield very high mortality (up to 24%), miscarriage and poor neonatal outcomes [1]; it may be speculated that these strains are yet to emerge in Nepal. Since there was no definitive evidence for this, a careful monitoring for drug resistance (including potential resistance genes) should be in place. Poor response with doxycycline in some countries suggested the potential emergence of resistant strains [25]. Early administration of doxycycline/azithromycin was found to reduce the progression to AES in India [3], treatment delay has to be avoided.

This study has some limitations. The detailed individual level data for was not available for many cases. IFA and PCR confirmation tests were limited to representative samples, and further genetic characterization pathogenic strain of *O. tsutsugamushi* that circulated in Nepal was not performed. Although a total of 104 rodent traps were set in the suspected areas, relatively small number of rodents were captured as well as low number of chigger mite was found. We cannot discount the possibility of other rodent species associated with scrub typhus in Nepal. Although we did not cover other rickettsial diseases, cases of spotted fever group, typhus group and Q Fever rickettsia in scrub typhus endemic areas of the neighbouring countries like India [18] and Bhutan [6] warrant further investigation of these etiologies in Nepal. Moreover, recent indication of Q fever in some acute undifferentiated febrile cases in Nepal also underscores it [21]. Apart from human, very high rickettsial seropositivity was reported among domestic animals in Bhutan suggesting the need of one health approach in controlling rickettsial diseases [34]. Therefore, the need of domestic animals studies also very relevant in context of Nepal. In conclusion, outbreak investigated in human, rodent and chigger mites with positive serological and molecular evidence, and an analysis of additional aggregated data confirm a firmly established scrub typhus infection ecology in Nepal. This is evidence of ongoing transmission of *O. tsutsugamushi* in Nepal. Persistently reported scrub typhus cases throughout the country for more than 2 years after the 2015 earthquake have imposed a risk for outbreaks of epidemic potential. Establishment of reference laboratory with IFA and molecular facility is urgently required in the country for confirmation, strain identification, genetic characterization and evolutionary analysis. Most importantly, the health system needs to be strengthened for systematic surveillance, early outbreaks detection and immediate response including treatment and preventive measures in the country.

## Acknowledgements

Authors thank Dr. Baburam Marasisni, then Director of Epidemiology and Diseases Control Division (EDCD), Department of Health Services, Dr Krishna Kumar Aryal, then Research Officer, Nepal Health Research Council for his contribution in research design and planning; the technical staffs, Mr. Resham Lamichhane, Mr. Uttam Raj Pyakurel, Mr.Bijay Rimal of EDCD, Department of Health Services, Mr. Shishir Pant of Vector Borne Disease Research and Training Centre (VBDRTC), Hetauda, Mr.Pramod Kumar Mehta from Shukraraj Tropical & Infectious Disease Hospital-Teku, Mr. Laxman Maharjan, and Mr. Buddha Ratna Maharajan of National Public Health Laboratory (NPHL) and Mr. Bijay Kumar Jha, Mr. Ram K.C. of District Public Health Office, Chitawan, Dr. Dayaram Lamsal, Dr. Santosh Pathak & Mr. Sanjay Yadav from CMC Hospital, Baratpur for their co-operation to conduct this study.

We are very grateful Dr. Wuttikon Rodk Vamtook, Mr. Surachai Leepita Krat and Mrs. Maneerat Somsri of Armed Forces Research Institute of Medical Sciences (AFRIMS), Thailand as WHO Collaborating Centre for Diagnostic Reference, Training and Investigation of Emerging Infectious Diseases for SEARO and Ms. Jasmine Shrestha, Mr. Pashupati Khanal, Mr. Ashish Shrestha, Mr. Bishnu Bahadur Rayamajhi, Ms. Samita Bajracharya & Ms. Bina Sakha of Walter Reed/ AFRIMS Research Unit Nepal (WARUN) for their technical support in confirmatory diagnosis of human samples and rodent/mite investigations.

## Supporting Information legends

**Supplementary Table 1: Distribution of Scrub typhus cases in in Nepal, 2015-2017**

